# Multiple approaches for massively parallel sequencing of HCoV-19 (SARS-CoV-2) genomes directly from clinical samples

**DOI:** 10.1101/2020.03.16.993584

**Authors:** Minfeng Xiao, Xiaoqing Liu, Jingkai Ji, Min Li, Jiandong Li, Lin Yang, Wanying Sun, Peidi Ren, Guifang Yang, Jincun Zhao, Tianzhu Liang, Huahui Ren, Tian Chen, Huanzi Zhong, Wenchen Song, Yanqun Wang, Ziqing Deng, Yanping Zhao, Zhihua Ou, Daxi Wang, Jielun Cai, Xinyi Cheng, Taiqing Feng, Honglong Wu, Yanping Gong, Huanming Yang, Jian Wang, Xun Xu, Shida Zhu, Fang Chen, Yanyan Zhang, Weijun Chen, Yimin Li, Junhua Li

## Abstract

COVID-19 has caused a major epidemic worldwide, however, much is yet to be known about the epidemiology and evolution of the virus. One reason is that the challenges underneath sequencing HCoV-19 directly from clinical samples have not been completely tackled. Here we illustrate the application of amplicon and hybrid capture (capture)-based sequencing, as well as ultra-high-throughput metatranscriptomic (meta) sequencing in retrieving complete genomes, inter-individual and intra-individual variations of HCoV-19 from clinical samples covering a range of sample types and viral load. We also examine and compare the bias, sensitivity, accuracy, and other characteristics of these approaches in a comprehensive manner. This is, to date, the first work systematically implements amplicon and capture approaches in sequencing HCoV-19, as well as the first comparative study across methods. Our work offers practical solutions for genome sequencing and analyses of HCoV-19 and other emerging viruses.

## INTRODUCTION

As of 14 March 2020, human coronavirus 2019 (HCoV-19) has surpassed severe acute respiratory syndrome coronavirus (SARS-CoV) and Middle East respiratory syndrome coronavirus (MERS-CoV) in every aspect, infecting over 140,000 people in more than 110 countries, with a mortality of over 5,000^1,2^. So far, coronaviruses have caused three major epidemics in the past two decades, posing a great challenge to global health and economy. Massively parallel sequencing (MPS) of viral genomes has demonstrated enormous capacity as a powerful tool to study emerging infectious diseases, such as SARS, MERS, Zika, and Ebola, in tracing the outbreak origin and drivers, tracking transmission chains, mapping the spread, and monitoring the evolution of the etiological agents^3–8^. Though by 14 March 2020, fewer than 500 HCoV-19 genomes were published on public databases including China National GeneBank DataBase (CNGBdb), NCBI GenBank, the Global initiative on sharing all influenza data (GISAID), etc, and much remains unknown about the epidemiology and evolution of the virus. One possible explanation of the paucity of published HCoV-19 genomes was the challenges posed by sequencing clinical samples with low virus abundance.

The first teams obtained the HCoV-19 genome sequences through metatranscriptomic MPS, supplemented by PCR and Sanger sequencing of a combination of bronchoalveolar-lavage fluid (BALF) and culture^9–11^ or from BALF directly^12,13^. Experience from studying SARS-CoV showed that BALF from the lower respiratory tract was an ideal sample type with higher viral load^14^. However, BALF was not routinely collected from every patient, and human airway epithelial (HAE) cell culture is very labor-intensive and time-consuming, taking four to six weeks^10,15^. The University of Hong Kong team managed to get the whole-genome sequences through metatranscriptomic sequencing with Oxford Nanopore platform supplemented by Sanger sequencing from both nasopharyngeal and sputum specimens after single-primer amplification^16^. The United States scientists published the whole-genome sequence using oropharyngeal and nasopharyngeal specimens through Sanger and meta-transcriptomic sequencing with both Illumina and MinIon^17^. To date, multiplex PCR-based or hybrid capture-based whole-genome sequencing of HCoV-19, as well as comparative studies between different approaches, have not been reported on peer-reviewed journals.

Besides inter-individual variations, dissecting intra-individual dynamics of viruses also largely promotes our understanding of viral-host interactions, viral evolution and transmission as demonstrated for Ebola, Zika, Influenza, etc^6,18-20^. The analyses of intra-individual single nucleotide variations (iSNVs) and its allele frequency have also contributed to antiviral therapy and drug resistance, e.g., to reveal highly conserved genes during the outbreak that potentially serve as ideal therapeutic targets^19,21^. However, it is a challenge to accurately detect iSNVs from clinical samples, especially when the samples are subjected to extra steps of enrichment and amplification.

Therefore, we aim to comprehensively compare the sensitivity, inter-individual (variant) and intra-individual (iSNV) accuracy, and other general features of different approaches by systematically utilizing ultra-high-throughput metatranscriptomic, hybrid capture-based, and amplicon-based sequencing approaches to obtain genomic information of HCoV-19 from serial dilutions of a cultured isolate and directly from clinical samples. We present a reasonable sequencing strategy that fits into different scenarios, and estimate the minimal amount of sequencing data necessary for downstream HCoV-19 genome analyses. Our study, to-gether with our tailor-made experimental workflows and bioinformatics pipelines, offers very practical solutions to facilitate the studies of HCoV-19 and other emerging viruses in the future and would promote extensive genomic sequencing and analyses of HCoV-19 and other emerging viruses, underpinning more comprehensive real-time virus surveillance and more efficient viral outbreaks managing.

## RESULTS

### Design of the comparative study

We sampled eight specimens from COVID-19 patients in February 2020, including throat swab, nasal swab, anal swab and sputum specimens, and the corresponding cycle threshold (Ct) value of HCoV-19 qRT-PCR ranges from 18 to 32 (Table 1). We initially tried to boost the coverage and depth of the viral genome by ultra-deep metatranscriptomic sequencing with an average sequencing amount of 1,607,264,105 paired-end reads (Table 1). Although we managed to obtain complete viral genome assemblies for each specimen, the sequencing depth varied across specimens. Only 0.002%-0.003% of the total reads were assigned to the HCoV-19 in three samples (GZMU0014, GZMU0030 and GZMU0031) with Ct between 29-32, resulting in inferior sequencing depth (less than 100X) (Table 1). Isolating viruses and enriching them in cell culture might improve the situation, but this requires high-standard laboratory settings and expertise apart from being time-consuming. Also, unwanted mutations that are not concordant with original clinical samples may be introduced during the culturing process.

**Table 1.**
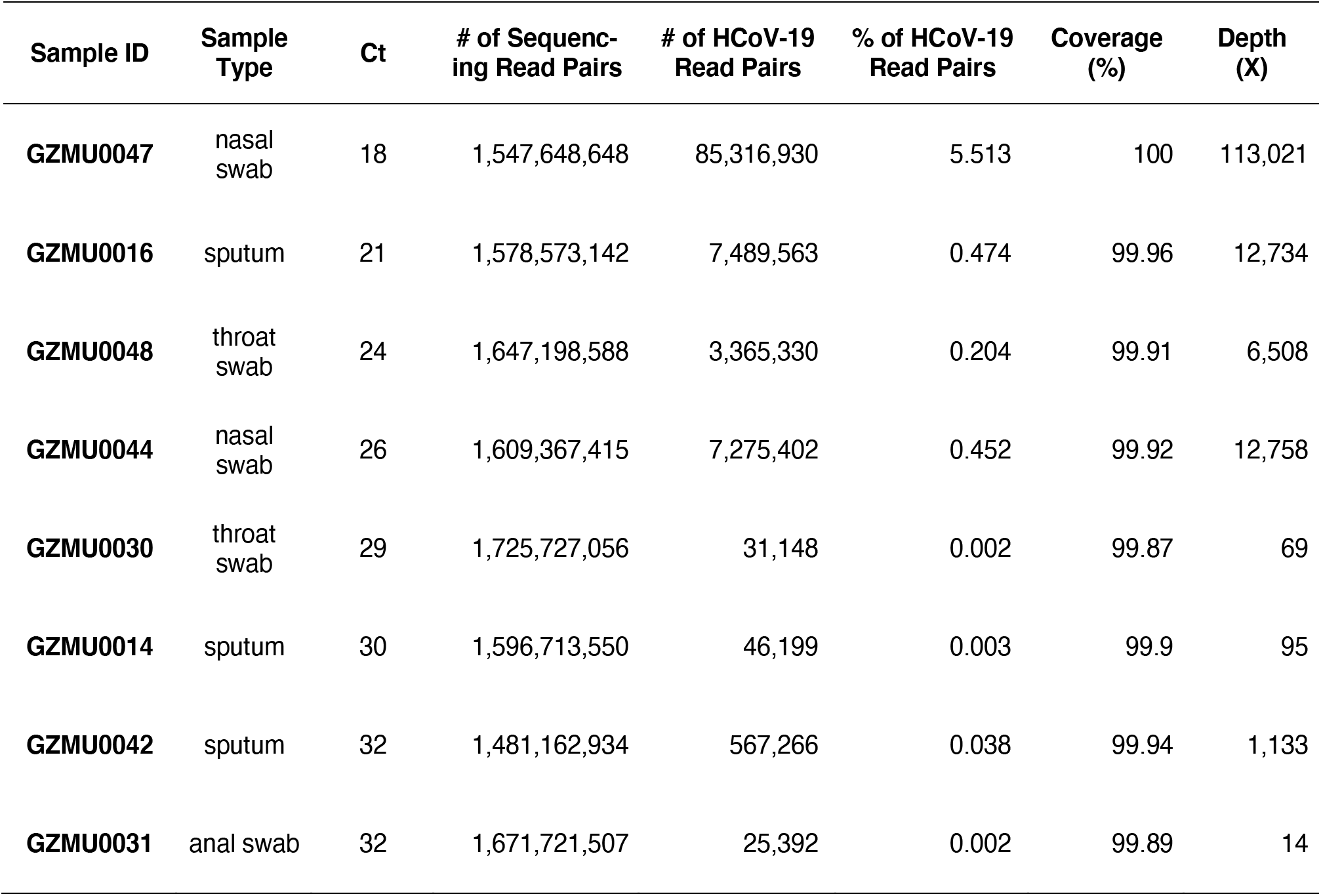
Metatranscriptomic sequencing data summary of eight HCoV-19 positive clinic al samples collected from Guangzhou in February 2020

To enrich adequate viral content for whole-genome sequencing in a convenient manner, we pursued two other methods: multiplex PCR amplification (amplicon) and hybrid capture (capture) (Fig. 1). We designed a systematic study to comprehensively validate the bias, sensitivity, inter-individual (variant) and intra-individual (iSNV) accuracy of multiple approaches by sequencing serial dilutions of a cultured isolate (unpublished), as well as the eight clinical samples (Fig. 2). We performed qRT-PCR of 10-fold serial dilutions (D1-D7) of the cultured isolate, and the Ct was 17.3, 20.8, 24.5 for, 28.7, 31.8, 35, and 39.9, respectively, indicating the undiluted RNA (D0) of the cultured isolate contained ~1E+08 genome copies per mL. For amplicon sequencing, we utilized two kits comprising of two set of primers generating PCR products of 300-400 bp and 100-200 bp, respectively. The ~400 bp amplicon-based sequencing was implemented in all samples and analyzed throughout the study, while the ~200 bp amplicon-based sequencing was only applied in the cultured isolate for coverage analysis.

**Figure 1.**
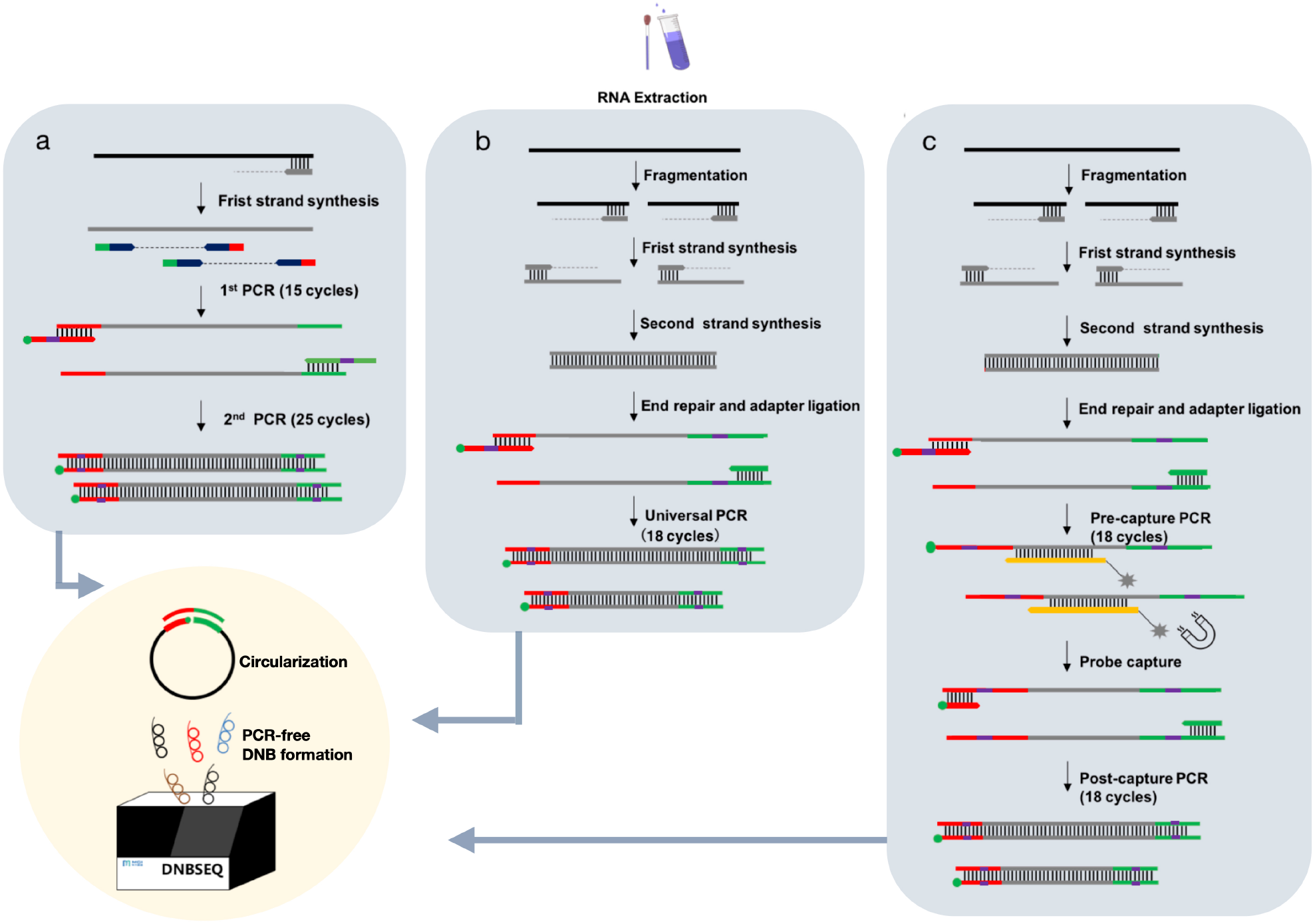
The general workflow of multiple sequencing approaches adopted in this study. We employed unique dual indexing (UDI) strategy and DNB-based (DNA Nanoball) PCR-free MPS platform to minimize index hopping and relevant sequencing errors^23,24,46^. **a**, Amplicon-based enrichment, the dual indexing was integrated in the 2nd PCR. Navy, multiplex PCR primers. **b**, Metatranscriptomic library preparations, the dual indexing was integrated in the universal PCR. **c**, Library preparations and hybrid capture-based enrichment, the 1st indexing was integrated in the pre-capture PCR while the 2nd indexing was integrated in the post-capture PCR. Ocher, ssDNA probes. Red and green lines represent adaptor sequences; green dots represent phosphate groups.

**Figure 2.**
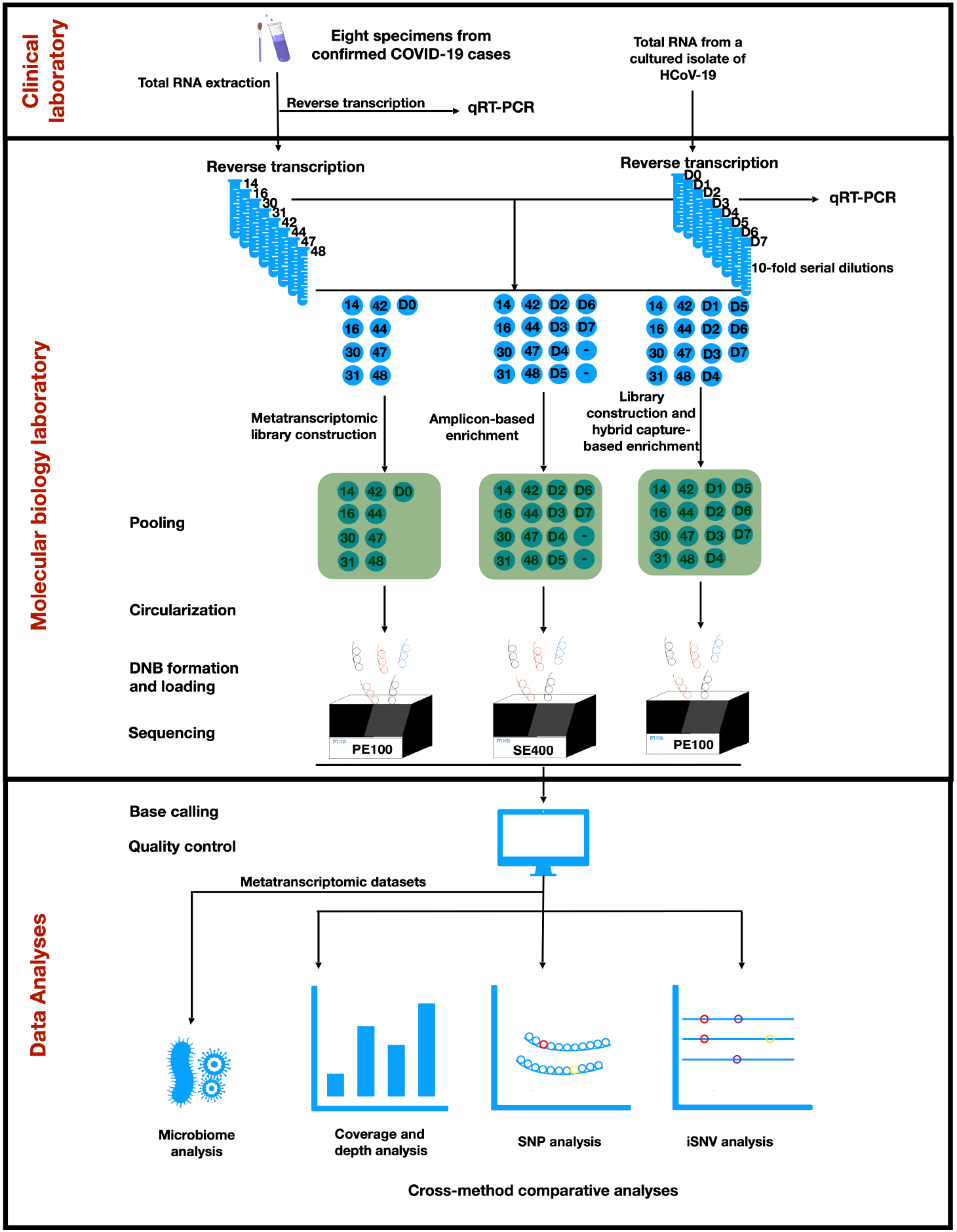
Overview of the study design. Eight clinical samples and serial dilutions of a cultured isolate were subjected to direct metatranscriptomic library construction, ampliconbased enrichment, and hybrid capture-based enrichment, respectively. Libraries generated from each method were pooled, respectively. DNB, DNA Nanoball. 14, GZMU0014; 16, GZMU0016; 30, GZMU0030; 31, GZMU0031; 42, GZMU0042; 44, GZMU0044; 47, GZMU0047; 48, GZMU0048. D0, undiluted sample of the cultured isolate; D1-D7, seven serial diluted samples of the cultured isolate, ranging from 1E+07 to 1E+01 genome copies per mL, in 10-fold dilution. -, negative controls prepared from nuclease-free water and human nucleic acids. PE100, paired-end 100-nt reads; SE400, single-end 100-nt reads.

### Comparison of evenness and sensitivity

Theoretically, amplicon sequencing should be the most sensitive and economical method among the three, and is particularly suitable in an outbreak where viral isolates are highly related. Although, there are still potential pitfalls, for instance, the 40 cycle-PCR in our workflow might augment trace amounts of HCoV-19 cross-contamination. To ensure the confidence of the datasets, we included serial dilutions of the cultured isolate and negative controls prepared from nuclease-free water and human nucleic acids since the 1st PCR. All samples in ~400 bp amplicon-based sequencing exhibited > 99.5% coverage across the HCoV-19 genome except for 1E+01 (95.23%), GZMU0031 (73.65%), HNA (6.17%), water (60.24%), suggesting the primers were well designed and the positive datasets were reliable. We also set stringent and method-specific criteria to filter low-confidence sequencing reads and samples (see Methods), e.g., clinical sample GZMU0031 was excluded for downstream sensitivity and accuracy validation due to inadequate depth in amplicon sequencing (Fig. 3a). Another pitfall is that amplification across the genome can hardly be unbiased, causing difficulties in complete genome assembly. Indeed, amplicon sequencing exhibited a higher level of bias compared with meta-transcriptomic sequencing, in terms of coverage across the viral genomes from the cultural isolate and the clinical samples tested in our study (Fig. 3b, d, Supplementary Fig. 1). To our surprise, however, capture sequencing was almost as unbiased as meta sequencing, demonstrating better performance than the previous capture method used to enrich ZIKV despite the HCoV-19 genome is ~3 fold larger than ZIKV^22^ (Fig. 3b, c). Two reasons amongst others were likely to be accountable to this improvement, 1) we utilized 506 pieces of 120 ssDNA probes covering 2x of the HCoV-19 genome to capture the libraries, 2) we employed the DNBSEQ sequencing technology that features PCR-free rolling circle replication (RCR) of DNA Nanoballs (DNBs)^23,24^.

**Figure 3.**
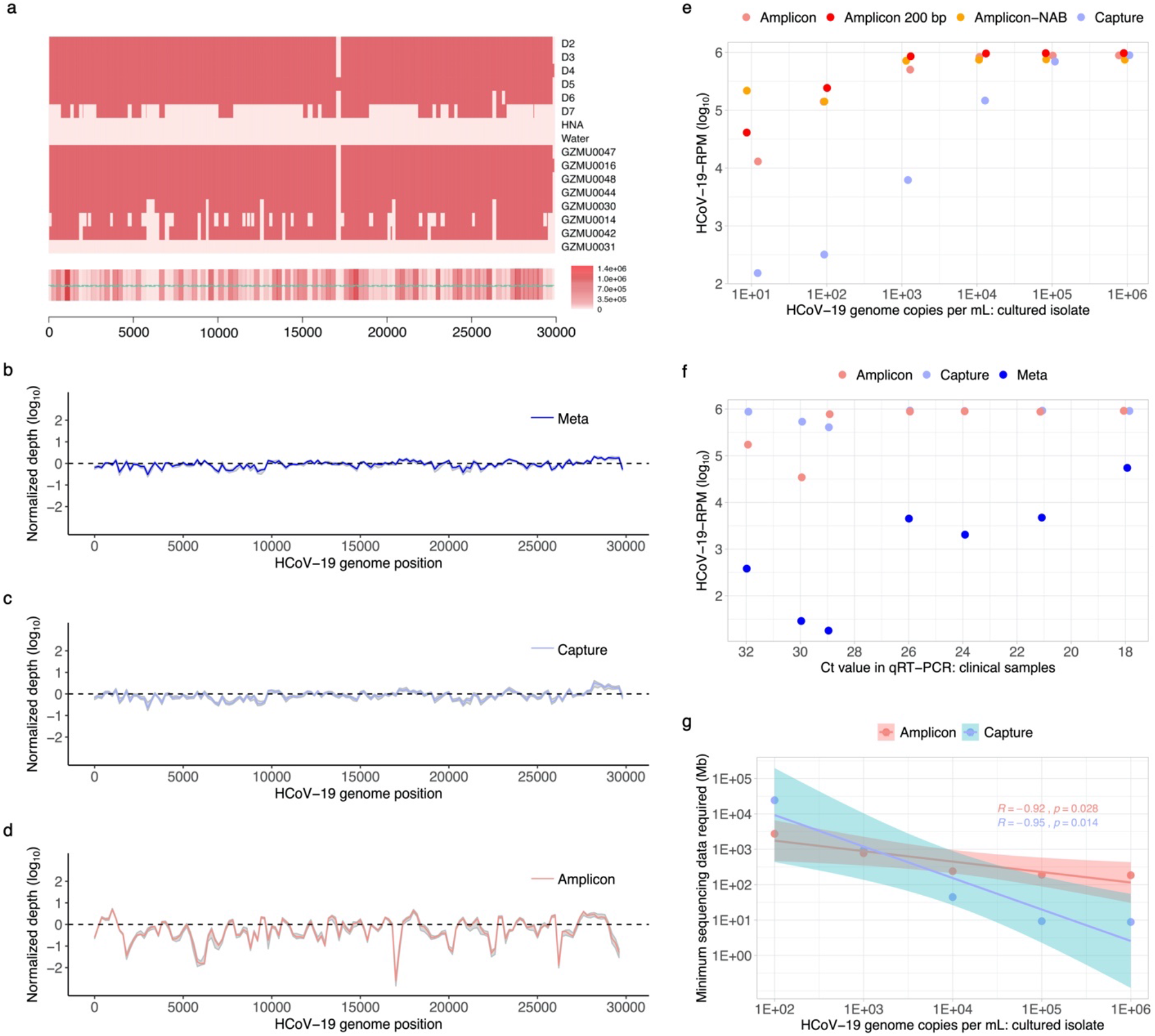
Sequencing coverage and depth of the cultured isolate and eight clinical samples. **a**, Amplicon sequencing coverage by sample (row) across the HCoV-19 genome. Pink, sequencing depth ≥100×; heatmap (bottom) sums coverage across all samples. HNA, negative control prepared from human nucleic acids; water, negative control prepared from nuclease-free water. Green horizontal lines on heatmap, amplicon locations. Overlap regions between amplicons range from 59-209 bp. **b-d**, Normalized coverage across viral genomes of the clinical samples across methods. **e**, HCoV-19-RPM sequenced plotted against genome copies per mL for the cultured isolate. Three independent experiments were performed for amplicon sequencing. Pink, ~400 bp amplicon-based sequencing including human and lambda phage nucleic acids background; red, ~200 bp amplicon-based sequencing; orange, ~400 bp amplicon-based sequencing excluding human and lambda phage nucleic acids background (NAB); light blue, capture sequencing. **f**, HCoV-19-RPM (Reads Per Million) sequenced plotted against qRT-PCR Ct value for the clinical samples. Pink, amplicon; light blue, capture; blue, meta. **g**, Estimated minimum amount of bases required by each method for high-confidence downstream analyses. Pink, amplicon; light blue, capture.

The sequencing results of amplicon and capture approaches revealed dramatic increases in the ratio of HCoV-19 reads out of the total reads compared with meta sequencing, suggesting the enrichment was highly efficient - 5596-fold in capture method and 5710-fold in amplicon method for each sample on average (Supplementary Table 1-2). To further compare the sensitivity of different methods, we plotted the number of HCoV-19 reads per million (HCoV-19-RPM) of total sequencing reads against the viral concentration for each sample. The productivity was similar between the two methods when the input RNA of the cultured isolate contained 1E+05 genome copies per mL and above (Fig. 3e). However, amplicon sequencing produced 10-100 fold more HCoV-19 reads than capture sequencing when the input RNA concentration of the cultured isolate was 1E+04 genome copies per mL and lower, suggesting amplicon-based enrichment was more efficient than capture for more challenging samples (conc. ≤ 1E+04 genome copies per mL, or Ct ≥ 28.7) (Fig. 3e). Meta sequencing - as expected - produced dramatically lower HCoV-19-RPM than the other two methods among clinical samples tested with a wide range of Ct values, whereas amplicon and capture were generally comparable to each other (Fig. 3e). Considering the costs for sequencing, storage, and analysis increase substantially with larger datasets, we tried to estimate how much sequencing data must be produced for each approach in order to achieve 10X depth across 95% of the HCoV-19 genome, and the results can be found in Supplementary Table 3. As a practical, cost-effective guidance for future sequencing, we also assessed the minimum amount of data required to pass the stringent filters (≥ 95% coverage and method-specific depth, see Methods) in our pipelines corresponding to different viral loads. We estimated that for high-confidence downstream analyses, amplicon sequencing requires at least 2,757 to 186 Mega bases (Mb) for samples containing 1E+02 to 1E+06 copies of HCoV-19 genome per mL, while capture sequencing requires 24,474 to 9 Mb for the same situation (Fig. 3g, Supplementary Table 4).

### Investigation of inter-and intra-individual variations

To determine the accuracy of different approaches in discovering inter-individual genetic diversity, we tested each method in calling the single nucleotide variations (SNVs) and verified some of the SNVs with Sanger sequencing (Supplementary Fig. 2). Two to five SNVs were identified within each clinical sample, and in all the seven samples, SNVs identified by the three methods were concordant except that capture missed one SNV at position 16535 in GZMU0014 (Fig. 4a). We then investigated the allele frequencies of these sites across methods, and found that alleles identified by capture sequencing displayed lower frequencies than the other two methods, especially for GZMU0014, GZMU0030, and GZMU0042 where the viral load was lower (Ct ? 29), which explained why capture sequencing neglected an SNV in our pipeline when the cutoff of SNV calling was set as 80% allele frequency (Fig. 4b). These data indicate that amplicon sequencing is more accurate than capture sequencing in identifying SNVs, especially for challenging samples.

**Figure 4.**
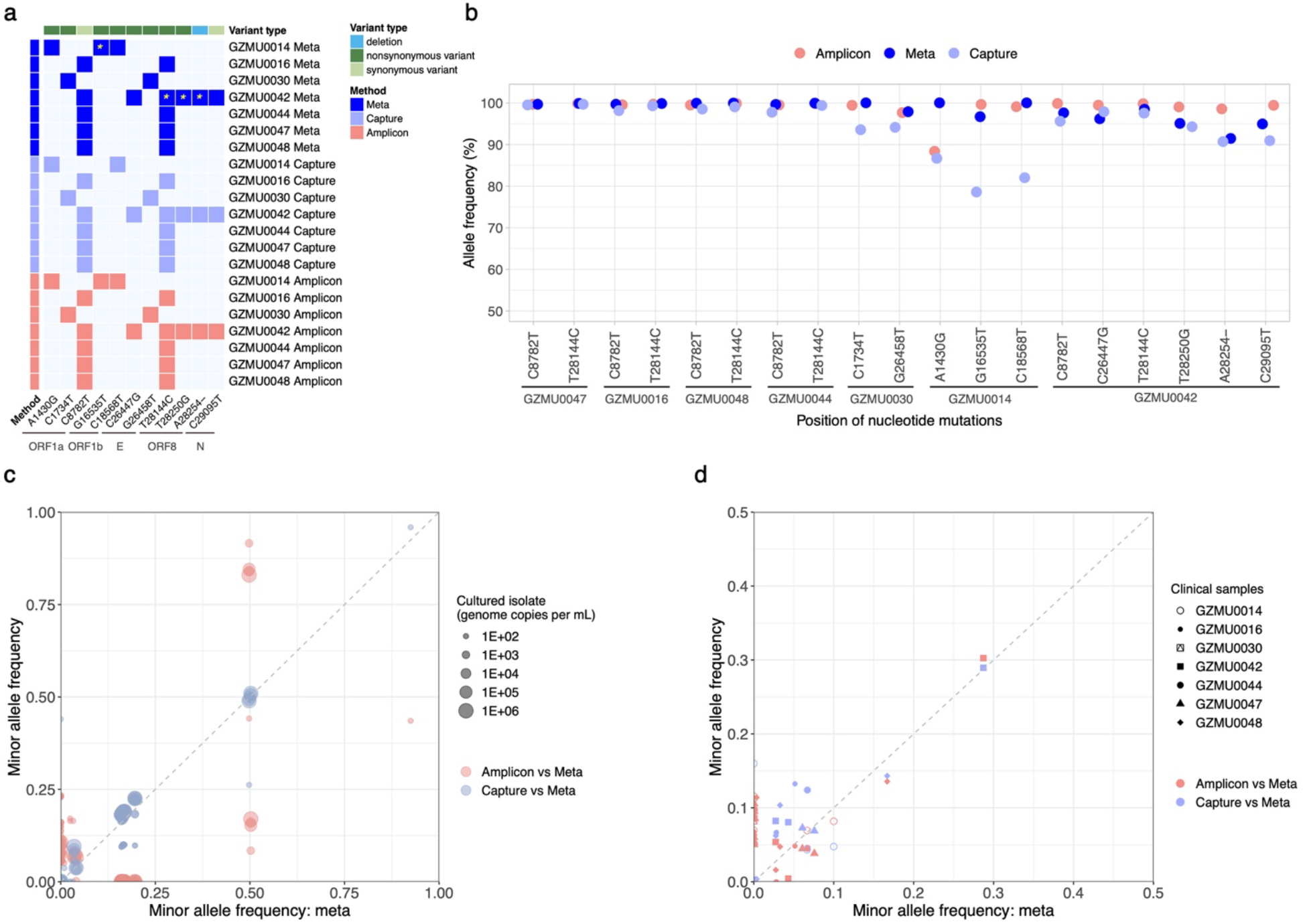
Between-sample and within-sample variants of HCoV-19 detected across methods. **a**, SNVs detected between clinical samples against a reference genome (GISAID accession: EPI_ISL_402119). Alleles with ≥ 80% frequencies were called. *, SNVs verified by Sanger sequencing. **b**, Allele frequencies of the identified SNVs. Pink, amplicon; light blue, capture; blue, meta. Minor allele frequencies detected in serial dilutions of the cultured isolate (**c**) and clinical samples (**d**) across methods. Pink, amplicon vs meta; light blue, capture vs meta. Minor alleles are defined with ≥ 5% and < 50% frequencies. Besides general quality filter, iSNVs had to pass depth and strand bias filter as described in Methods.

To further determine the accuracy of different approaches in identifying HCoV-19 iSNVs, we examined minor allele frequencies in serial dilutions of the cultured HCoV-19 isolate and clinical samples. For serial dilutions of the cultured isolate, the minor allele frequencies detected in capture sequencing datasets were generally approximate to meta sequencing, while most allele frequencies in amplicon sequencing datasets deviated with those in meta sequencing (Fig. 4c) A similar pattern was shown for clinical samples, indicating that amplicon sequencing was unreliable of quantifying minor allele frequencies (Fig. 4d). Plotting allele frequencies against HCoV-19 concentrations supported the above finding, and further revealed that amplicon sequencing was unreliable of allele frequencies at all concentrations while capture sequencing was reliable at > 1E+03 genome copies per mL (Supplementary Fig. 3). Referring to the iSNV identified in clinical samples by meta sequencing, we then calculated the false positive rate (FPR) and false negative rate (FNR) of minor alleles called by amplicon and capture methods. The FPR and FNR of minor alleles identified in amplicon sequencing was 0.74% and 66.67%, while that in capture sequencing was 0.02% and 0%, respectively. Together these results suggest amplicon sequencing was not as accurate as capture sequencing in identifying minor alleles, which could be in part due to Matthew effect caused by PCR.

### Microbiome in clinical samples

In addition to target viral genome, metatranscriptomic sequencing has also allowed us to investigate RNA expression patterns of the overall microbiome and host content and thus suitable for discovering new viruses, distinguishing coinfections, and dissecting virus-host interactions. To explore the microbiota, we performed further metatranscriptomics analysis of the clinical samples. We were able to identify host nucleic acids in all of the samples, and over 95% of total reads were from the host in sputum, nasal, and throat samples (Supplementary Fig. 4a). Virus contributed to less than 5% of reads in anal swab and throat swab while more than 50% of reads in nasal swab (Supplementary Fig. 4b). These results suggest nasal swab could be the most ideal sample type for viral detection among the four sample types, which agrees with recent clinical evidence^25^. Among the viral reads, over 90% were Coronaviridae, which is consistent with clinical diagnostics (Supplementary Fig. 4c). Reads from other viruses were also identified, indicating further measurements could be taken to confirm if co-infection exists (Supplementary Fig. 4). Bacterial composition was also shown, providing support for scientific research, as well as for further confirmation of bacterial infection and antibiotics prescription (Supplementary Fig. 4d-f).

### Guidance for virus sequencing

Taken together, each sequencing scheme elaborated here for massively parallel sequencing of HCoV-19 genomes has its own merits (Table 2). We hereby propose a reasonable, cost-effective strategy for sequencing and analyzing HCoV-19 under different situations: 1) if one wants to study other genetic materials than the target viruses, or the viruses become highly diversified via recombinational events, or the viral load within the RNA sample is high (e.g. conc. ≥ 1E+05 viral genome copies per mL, or Ct ≤ 24.5), meta sequencing can be prioritized; 2) if one focuses on intra-individual variations for more challenging samples (e.g. conc. < 1 E+05 viral genome copies per mL, or Ct > 24.5), capture sequencing seems to be a justified choice; and, 3) if identifying SNVs is the main purpose, the most convenient, economical strategy would be amplicon sequencing that can support high-confidence analyses of samples containing viral content as low as 1E+02 viral genome copies per mL, or Ct as high as 35.

**Table 2.** General characteristics of the three approaches employed in this study

|  | Metatranscriptomic sequencing | Hybrid capture-based sequencing | Multiplex PCR amplicon-based sequencing |
| --- | --- | --- | --- |
| Sequencing objective | Microbiome+Human | Target genome | Target genome |
| 2nd strand synthesis | Y | Y | N |
| Fragmentation | Y | Y | N |
| Library preparation | Y | Y | N |
| PCR | 18 cycles | 18+18 cycles | 15+25 cycles |
| Estimated time for pre-sequencing sample processing | 10.5 h | 20.5 h | 7.5 h |
| Oligo synthesis | - | 120 nt x 506 | 40-60 nt x 2 x (113+14+10) |
| Cost estimated for pre-sequencing sample processing | Moderate | High | Low |
| Estimated minimum data for downstream analyses (Base level) | >10Gb | Mb | Mb |
| Evenness | High | Moderate | Low |
| Sensitivity | + | ++ | +++ |
| Accuracy (SNV) | +++ | ++ | +++ |
| Accuracy (iSNV) | +++ | ++ | + |

## DISCUSSION

Sequencing low-titre viruses directly from clinical samples is challenging, which is further exacerbated by the fact that coronavirus genomes are the largest among RNA viruses (~3 fold larger compared with ZIKV). Compared with direct metatranscriptomic sequencing, high sensitivity of hybrid capture and amplicon sequencing methods come at a price of low accuracy, and neither of the two can be used to sequence highly diverse or recombinant viruses because the primers and probes are specific to known viral genomes. Amplicon sequencing compromises its accuracy, while it becomes the most convenient and economical method of all. Either or a combination of the approaches described here can be chosen to cope with various needs of researchers, e.g., metatranscriptomic sequencing data with insufficient coverage and depth can be pooled with hybrid capture data to generate high quality assemblies^22^. Our research, as well as the methods elaborated here, are able to help other researchers to sequence and analyze large viruses from clinical samples and thus benefit investigations on the genomic epidemiology of viruses.

Some pros and cons described above might be specific to the experimental workflows and bioinformatics pipelines tailored in this study, e.g., 1) the bias of amplicon sequencing can be improved by reducing the amount of cycles in the 1st PCR or optimize the molar ratios of primers (Fig. 1a), 2) the amplicon sequencing is particularly convenient compared with previous counterparts since the fragmentation and library construction steps are omitted here by integrating adaptor and barcode ligation in the 2nd PCR and sequencing the amplicons using single-end 400 nt reads (Fig. 1a), 3) using anything less than 506 pieces of 120 ssDNA probes in hybrid capture may attenuate the sequencing coverage while increase the bias, 4) metatranscriptomic sequencing was conducted with an ultra-high-throughput sequencing platform so that the successful rate was substantially higher than usual. 5) the minimal amount of data necessary for analyzing the HCoV-19 genome from clinical samples across methods is higher than that predicted by data from the cultured isolate was probably due to the high nucleic acids background from the host and other microbes (Supplementary Table 3-4, Supplementary Fig. 4). Also, we do not take into account the time spent in sequencing since the workflows can be easily adapted in order to be compatible with various platforms including Illumina and Oxford Nanopore Technologies (ONT), besides DNBSEQ of MGI.

## METHODS

### Ethics statement

The Institutional Review Boards (IRB) of the First Affiliated Hospital of Guangzhou Medical University approved the clinical studies. IRB of BGI-Shenzhen approved the sequencing and downstream studies.

### Sampling, RNA extraction, reverse transcription and qRT-PCR

Clinical specimens (including throat swab, nasal swab, anal swab, and sputum) were obtained from confirmed COVID-19 cases at the First Affiliated Hospital of Guangzhou Medical University. Total RNA of the cultured isolate of HCoV-19 was obtained from the Academy of Military Medical Science (AMMS), and subjected to 10-fold serial dilutions. Total RNA was extracted with QiAamp RNeasy Mini Kit (Qiagen, Heiden, Germany) following the manufacturer’s instructions without modification. Real-time reverse transcription PCR (qRT-PCR) targeting RdRp gene and N gene of HCoV-19 was used to detect and quantify the viral RNA within clinical samples and serial dilutions of the cultured isolate using the HCoV-19 Nucleic Acid Detection Kit following the manufacture’s protocol (Geneodx, Shanghai, China, and BGI-Shenzhen, Shenzhen, China).

### Metatranscriptomic library preparation and sequencing

Host DNA was removed from RNA samples using DNase I, and the concentration of RNA samples was measured by Qubit RNA HS Assay Kit (Thermo Fisher Scientific, Waltham, MA, USA). DNA-depleted and purified RNA was used to construct the single-stranded circular DNA library with MGIEasy RNA Library preparation reagent set (MGI, Shenzhen, China), as follows: 1) RNA was fragmented by incubating with fragmentation buffer at 87°C for 6 min; 2) double-stranded (ds) cDNA was synthesized using random hexamers with fragmented RNA; 3) ds cDNA was subjected to end repair, adaptor ligation, and 18-cycle PCR amplification; 4) PCR products were Unique Dual Indexed (UDI), before going through circularization, and rolling circle replication (RCR) to generate DNA nanoballs (DNBs)-based libraries. DNBs preps of clinical samples were sequenced on the ultra-high-throughput DNB-SEQ-T7 platform (MGI, Shenzhen, China) with paired-end 100 nt strategy, generating 321 Gb sequencing data for each sample on average.

### Hybrid capture-based enrichment and sequencing

A hybrid capture technique was used to enrich HCoV-19-specific content from the meta-transcriptomic double-stranded DNA libraries with the 2019-nCoVirus DNA/RNA Capture Panel (BOKE, Jiangsu, China). Manufacturer’s instructions were slightly modified to accommodate the MGISEQ-2000 platform, i.e., blocker oligos and PCR primer oligos were replaced by MGIEasy exon capture assistive kit (MGI, Shenzhen, China). DNBs-based libraries were constructed and sequenced on the MGISEQ-2000 platform with paired-end 100 nt strategy using the same protocol described above, generating 37 Gb sequencing data for each sample on average.

### Amplicon-based enrichment and sequencing

Total RNA was reverse transcribed to synthesize the first-strand cDNA with random hexamers and Superscript II reverse transcriptase kit (Invitrogen, Carlsbad, USA). Sequencing was attempted on all samples regardless of Ct value including negative controls prepared from nuclease-free water and NA12878 human gDNA. A two-step HCoV-19 genome amplification was performed with an equimolar mixture of primers using ATOPlex SARS-CoV-2 Full Length Genome Panel following the manufacture’s protocol (MGI, Shenzhen, China), generating 137X ~400 bp amplicons or 299X ~200 bp amplicons and the genome positions of the amplicons are shown in Supplementary Table 5. 20 μl of first-strand cDNA was mixed with the components of the first PCR reaction following the manufacturer’s instructions, including lambda phage gDNA unless specified. 2 ng of Human gDNA was added to each PCR reaction of the cultured isolate. The PCR was performed as follows: 5 min at 37°C, 10 min at 95°C, 15 cycles of (10 s at 95°C, 1 min at 64°C, 1min at 60°C to 10 s at 72°C), 2 min at 72°C. The products were purified with MGI EasyDNA Clean beads (MGI, BGI-Shenzhen, China) at a 5:4 ratio and cleaned with 80% concentration ethanol according to the manufacturer’s instructions. The 2nd PCR was performed under the same regimen as the 1st PCR except for 25 cycles, and the beads-purified products from the first PCR reaction were unique dual indexed. After the 2nd PCR, products were purified following the same procedures as the 1st PCR and quantified using the Qubit dsDNA High Sensitivity assay on Qubit 3.0 (Life Technologies). PCR products of samples yielding sufficient material (> 5 ng/μl) were pooled at equimolar to a total DNA amount of 300 ng before converting to single-stranded circular DNA. DNBs-based libraries were generated from 20 μl of single-stranded circular DNA pools and sequenced on the MGISEQ-2000 platform with single-end 400 nt strategy, generating 1.8 Gb sequencing data for each sample on average.

### Identification of HCoV-19-like reads from Massively Parallel Sequencing data

For metatranscriptomic and hybrid capture sequencing data, total reads were first processed by Kraken v0.10.5^26^ (default parameters) with a self-build database of Coronaviridae genomes (including SARS, MERS and HCoV-19 genome sequences downloaded from GISAID, NCBI and CNGB) to efficiently identify candidate viral reads with a loose manner. These candidate reads were further qualified with fastp v0.19.5^27^ (parameters: -q 20 -u 20 - n 1 -l 50) and SOAPnuke v1.5.6^28^ (parameters: -l 20 -q 0.2 -E 50 -n 0.02 -5 0 -Q 2 -G -d) to remove low-quality reads, duplications and adaptor contaminations. Low-complexity reads were next filtered by PRINSEQ v0.20.4^29^ (parameters: -lc_method dust -lc_threshold 7). After the above process, HCoV-19-like reads generated from metatranscriptomics and hybrid capture method were obtained.

For amplicon sequencing data, SE 400 reads were first processed with fastp v0.19.5^27^ (parameters: -q 20 -u 20 -n 1 -l 50) to remove low quality-reads and adaptor sequences. Primer sequences and the 21 nt upstream and downstream of primers within the reads were then trimmed with BAMClipper v1.1.1^30^ (Parameters: -n 4 -u 21 -d 21). Reads with low quality bases, adaptors, primers and adjacent sequences completely removed as described above were considered as clean reads for downstream analyses.

### Assembling viral genome

HCoV-19-like reads of metatranscriptomic and hybrid capture sequencing data were *de novo* assembled with SPAdes (v3.14.0)^31^ using the default settings to obtain virus genome sequences. To reduce the complexity of the assembling process, identified HCoV-19-like reads of metatranscriptomic and hybrid capture sequencing data were subsampled to the data amount greater than 100X sequencing depth for the HCoV-19 genome. For the two metatranscriptomic samples with a sequencing depth lower than 100X for the HCoV-19 genome (GZMU0014 and GZMU0030), all HCoV-19-like reads were used for assembling viral genomes.

Due to the uneven read coverage in amplicon sequencing of HCoV-19, virus consensus sequences of amplicon samples were generated by Pilon v1.23^32^ (parameters: --changes - vcf --changes --vcf --mindepth 1 --fix all, amb). Positions with depth less than 100X or lower five times than negative control samples were masked as ambiguous base N.

### Assessment the coverage depth across the viral genome

HCoV-19-like reads of metatranscriptomic and hybrid capture sequencing data were aligned to the HCoV-19 reference genome (GISAID accession: EPI_ISL_402119) with BWA aln (v0.7.16)^33^. Duplications were identified by Picard Markduplicates (v2.10.10)(http://broadinstitute.github.io/picard) with default settings. For each sample, we calculated the depth of coverage at each nucleotide position of the HCoV-19 reference genome with Samtools (v1.9)^34^ and scaled the values to the mean depth. For each nucleotide position, we calculated the median depth, and 20th and 80th percentiles across all samples. Read coverage and depth across the HCoV-19 reference genome were plotted by a 200-nt sliding window with the ggplot2^35^ package in R (v3.6.1)^36^.

Amplicon sequencing data were processed as described above, except that duplications were not removed. A heatmap was generated to visualize the viral genome coverage for all samples sequenced by the amplicon method with the pheatmap^37^ package in R (v3.6.1)^36^. The depth at each nucleotide position was binarized, and was shown in pink if the depth was 100x and above.

### Relationships between genome copies and method-dependent minimum amount of sequencing data

HCoV-19 reads of metatranscriptomic and hybrid capture sequencing data were identified by aligning the HcoV-19-like reads to the HCoV-19 reference genome (GISAID accession: EPI_ISL_402119) with BWA in a strict manner of coverage ≥ 95% and identity ≥ 90%. For comparisons of the coverage and depth of the viral genome across samples and methods, we normalized the viral reads to total sequencing reads with HCoV-19 Reads Per Million (HCoV-19-RPM). HCoV-19-RPM for amplicon sequencing data was calculated by the same pipelines we applied for metatranscriptomic and hybrid capture sequencing data.

To estimate the minimum data requirements for genome assembling and intra-individual variation analysis, we applied gradient-based sampling to the HCoV-19 genome alignments (referred to BAM files) to each dataset using Samtools (v1.9)^34^. The effective genome coverage was set as 95% for all three MPS methods. Considering the distinct technologies used in different methods, we set method-dependent thresholds of effective depth as follows: 1) ≥ 10X for metatranscriptomic sequencing; 2) ≥ 20X for hybrid capture sequencing; and 3) ≥ 100X for amplicon sequencing. We next calculated the coverage and depth within each subsampled BAM file per sample to determine the minimal BAM file that could meet the above thresholds of both coverage and sequencing depth. The methoddependent minimum amount of sequencing data of each sample were estimated accordingly. We assessed the correlations between the HCoV-19 genome copies per mL in diluted samples of cultured isolates and the minimum amount of sequencing data for amplicon-and capture-based methods using Pearson correlation coefficient (R) with the function *scatter* from the R package *ggpubr* (v3.6.1)^38^.

### Consistency in variants calling performance among methods

Except for amplicon sequencing samples, variants calling in metatranscriptomic and hybrid capture sequencing samples was performed in the previous BAM files of identified HCoV-19 reads after removing duplications from alignment output by Picard Markduplicates (http://broadinstitute.github.io/picard). To accurately identify SNVs from viral sequencing data of all three methods, we first called SNVs with freebayes (v1.3.1)^39^ (parameters: -p 1 - q 20 -m 60 --min-coverage 10 -V) and then filtered the low-confidence SNVs with snippy-vcf_filter^40^ (parameters: --minqual 100 --mincov 10 --minfrac 0.8). Remaining SNVs post filtering in VCF files generated by freebayes were annotated in HCoV-19 genome assemblies and consensus sequences with SNVeff (v4.3)^41^ using default parameters.

Next, we calculated SNV allele frequencies and called iSNVs (intra-host Single Nucleotide Variations) for each dataset to assess the consistency of variants calling performance among three methods. We performed pysamstats v1.1.2 (https://github.com/ali-manfoo/pysamstats) (parameters: -type variation_strand --min-baseq 20 -D 1000000) to count the number of matches, mismatches, deletions and insertions at each base, estimate nucleotide percentage and determine allele frequencies of SNVs at reference genome positions based on the HCoV-19 alignments from BAM files.

For amplicon sequencing data, only base positions covered by ≥100X reads were used for iSNVs calling. For metatranscriptomic and hybrid capture-based sequencing data, the thresholds of depth were set as 10X and 20X, respectively. The candidate iSNVs were further filtered for quality as follows: 1) frequency filtering, only minor alleles (frequency ≥ 5% and <50%) and major alleles (frequency ≥ 50% and ≤ 95%) were remained; 2) depth filtering, iSNVs with fewer than five forward or reverse reads were removed; and 3) strand bias filtering (not applicable to single-end reads of amplicon sequencing), iSNVs were removed if there were more than a 10-fold strand bias or a 5-fold difference between the strand bias of the variant call and that of the reference call.

### Taxonomy of clinical samples by unbiased metatranscriptomic sequencing

For metatranscriptomic sequencing of clinical samples, raw sequencing data of a single sequence lane (approximately 60-75 Gb per sample) was used to simultaneously assess the RNA expression patterns of human, bacteria and viruses in clinical samples from COVID-19 patients. We first used software fastp (v0.19.5)^27^ to filter low-quality reads and remove adapter with parameters: -5 -3 -q 20 -c -l 30. After QC, we mapped high-quality reads to hg19 and removed human ribosomal RNA (rRNA) reads by SOAP2 v2.21^42^ (parameters: - m 0 -x 1000 -s 28 -l 32 -v 5 -r 1), and the remaining RNA reads were then aligned to hg19 by HISAT2^43^ with default settings to identify non-rRNA human transcripts as previously described. Next, we applied Kraken 2^44^ (version 2.0.8-beta, parameters: --threads 24 --confidence 0) to assign microbial taxonomic ranks to non-human RNA reads against the large reference database MiniKraken2 (April 2019, 8GB) built from the Refseq bacteria, archaea, and viral libraries and the h38 human genome. Bracken^45^ (Bayesian Reestimation of Abundance with Kraken) was further applied to estimate microbial relative abundances based on taxonomic ranks of reads assigned by Kraken2.

## Supporting information

Supplementary Information combined

Supplementary Information combined

Supplementary Information combined

## Data availability

Sequencing data that support the findings of this study have been deposited in CNGB (https://db.cngb.org/) under Project accession CNP0000951 and CNP0000955, and in GISAID under accession EPI_ISL_414663, EPI_ISL_414686 to EPI_ISL_414692.

## Code availability

The software and parameters used in data analysis can be found in Supplementary Table 6.

## ACKNOWLEDGEMENTS

We attribute this work to the amazing people in this land who dedicate themselves to the battle of mankind against viruses. This work is funded by the State Key Research Development Program of China (2019YFC1200501), the National Major Project for Control and Prevention of Infectious Disease in China (2018ZX10301101-004), the emergency grants for prevention and control of SARS-CoV-2 of Ministry of Science and Technology (2020YFC0841400) and Guangdong province, China (2020B111107001, 2020B111108001, 2018B020207013), Guangdong Provincial Key Laboratory of Genome Read and Write (No. 2017B030301011), Guangdong Provincial Academician Workstation of BGI Synthetic Genomics (No. 2017B090904014), and Shenzhen Engineering Laboratory for Innovative Molecular Diagnostics (DRC-SZ[2016]884). We thank China National GeneBank at Shenzhen for providing sequencing service. We appreciate all the authors who have deposited and shared genome data on GISAID, and a table with genome sequence acknowledgments can be found in Supplementary Table 7.

## AUTHOR CONTRIBUTIONS

J.L., W.C. and M.X. conceived the project. X.L., J.Z, Y.W., and Y.L. sampled and processed the clinical specimen. M.X., Ji.L., M.L., and J.L. designed the experiments. L.Y. and Y.Z. developed the multiplex PCR amplicon-based sequencing method. M.L., Ji.L., Y.L, P.R. W.S., G.Y. and T.C. performed multiplex PCR and amplicon sequencing. J.L., and P.R. performed metatranscriptomic library construction and hybrid capture experiments. J.J., M.L, W.S., T.L., H.R., and H.Z. processed the sequencing data and conducted bioinformatic analyses. J.L., M.X. H.Z., J.J., M.L., and W.S. interpreted the data. M.X., J.J., M.L., and J.L. wrote and polished the manuscript. H.Z., W.S., L.Y., W.C. and Y.Z. contributed substantially to the manuscript revisions. All other authors provided useful suggestions and comments on the project and the manuscript.

## COMPETING INTERESTS

ATOPlex SARS-CoV-2 Full Length Genome Panel is a proprietary product.

PCR PRIMER PAIR AND APPLICATION THEREOF

Patent applicant: MGI Tech Co.,Ltd

Name of inventor(s): Lin Yang, Ya Gao, Guodong Huang, Yicong Wang, Yuqian wang, Yanyan Zhang, Fang Chen, Na Zhong, Hui Jiang, Xun Xu.

Application number: PCT/CN2017/089195

Any inquires or requests regarding this product should be specifically addressed to Yan-yan Zhang.

