## Supplementary Information combined for "Multiple approaches for massively parallel sequencing of HCoV-19 (SARS-CoV-2) genomes directly from clinical samples"

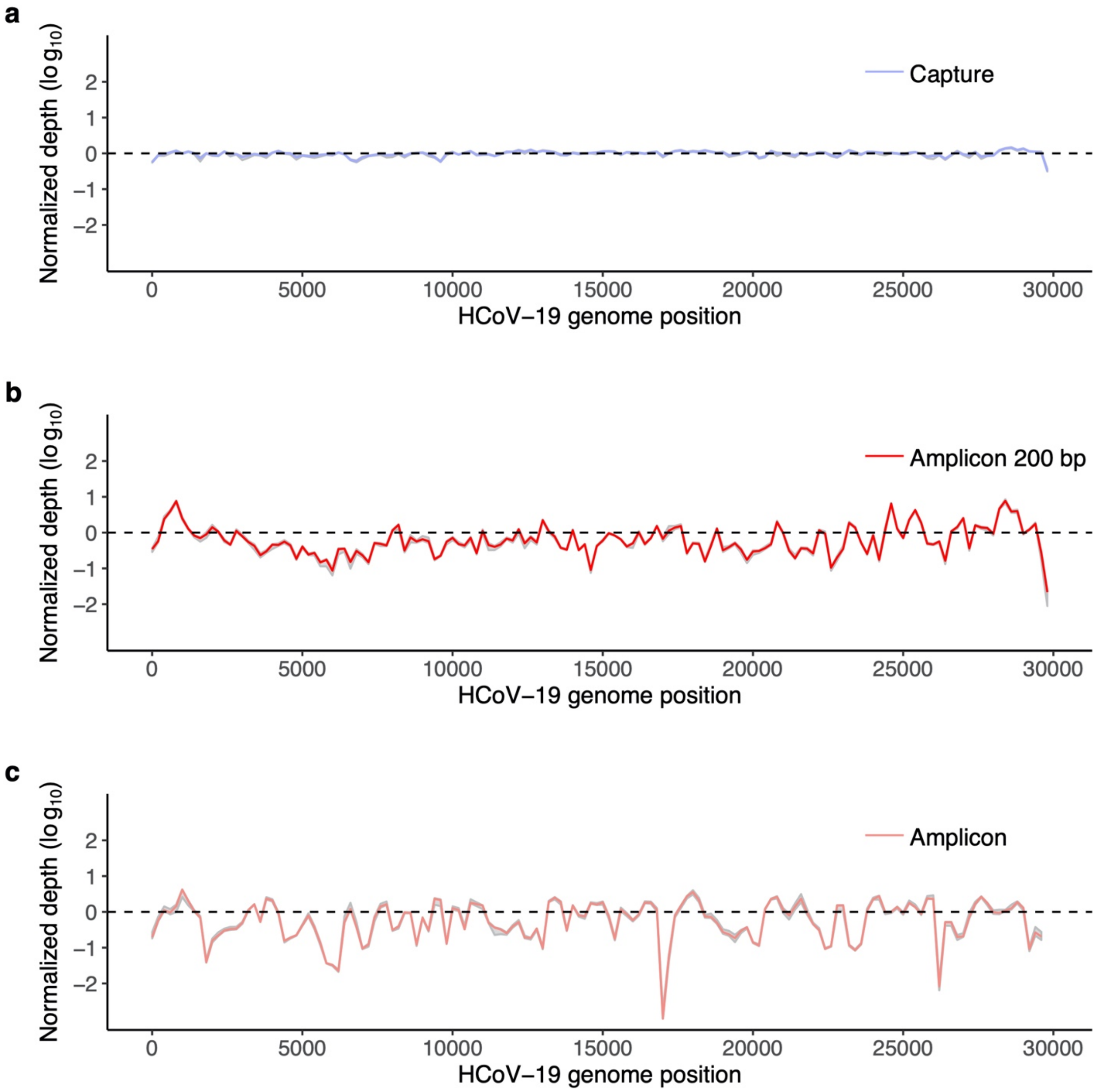


**Supplementary Figure 1. Normalized coverage across HCoV-19 genomes of serial dilutions of the cultured isolate sequenced by multiple approaches.** (**a**) Hybrid capture sequencing; (**b**) ~200 bp amplicon sequencing; (**c**) ~400 bp amplicon sequencing. Four dilutions of the cultured isolate (D2-D5, see details in Supplementary table 1) are used**.** The normalized depth at 50th (median, blue in **a**, red in **b**, and light red in **c**), and 20th and 80th percentiles (gray in a-c) of each position are shown.

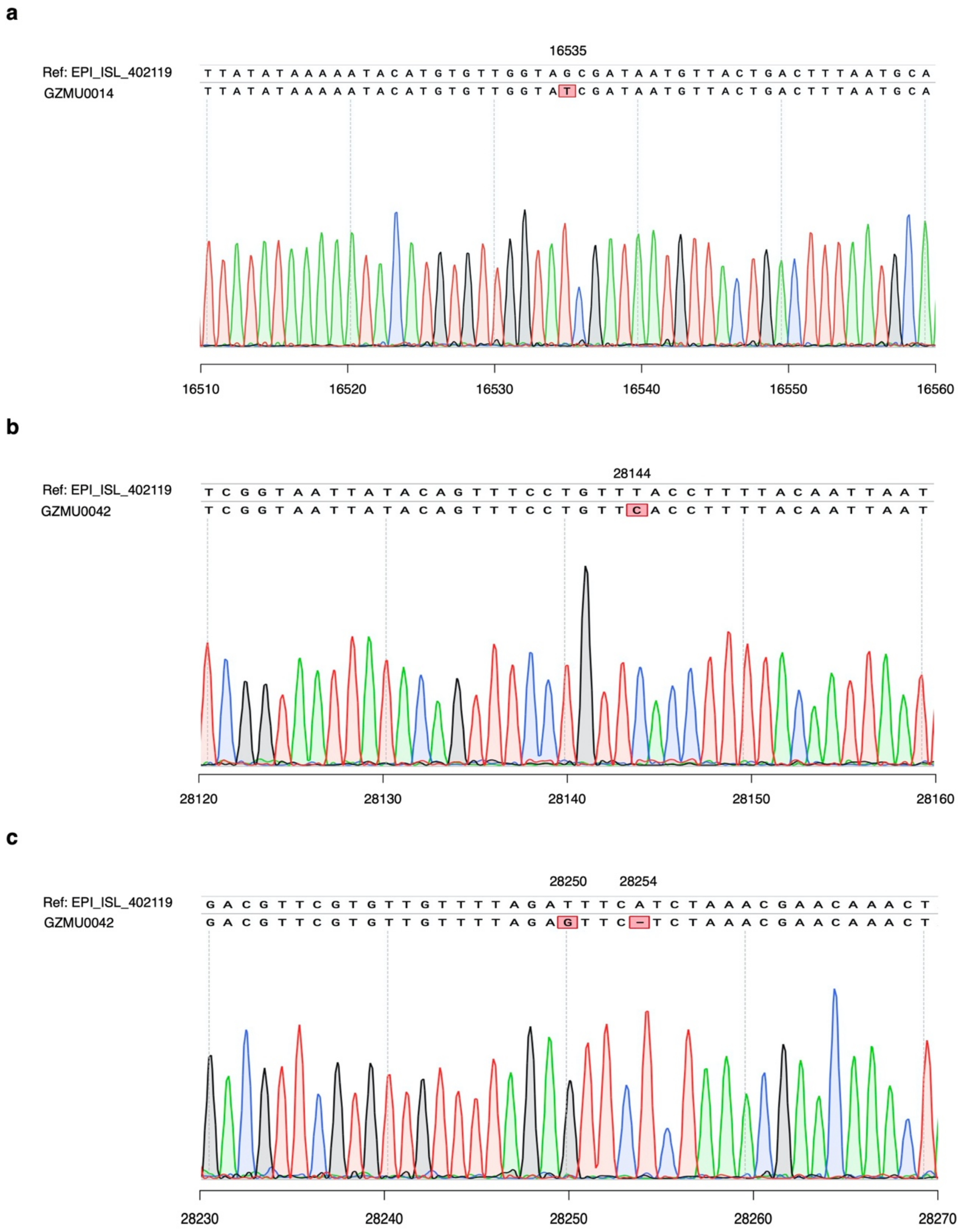


**Supplementary Figure 2. Sanger sequencing results of SNVs in clinical samples.** Shown are (**a**) G16535T mutation in GZMU0014, (**b**) T28144C mutation, (**c**) T28250G mutation and A28254- deletion in GZMU0042 aligned against the reference genome (GISAID accession: EPI_ISL_402119).

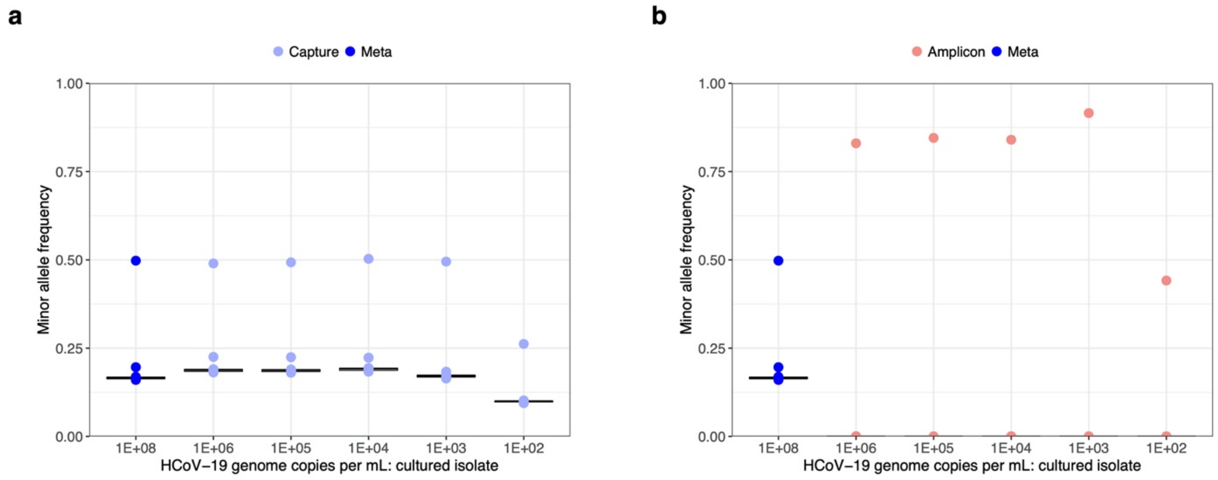


**Supplementary Figure 3. Box-plot of allele frequencies against HCoV-19 genome copies per mL.** Sixteen minor alleles of HCoV-19 cultured isolate were identified by metatranscriptomic sequencing (Meta, dark blue), compared with their allele frequencies identified by capture sequencing (**a**), and amplicon sequencing (**b**).

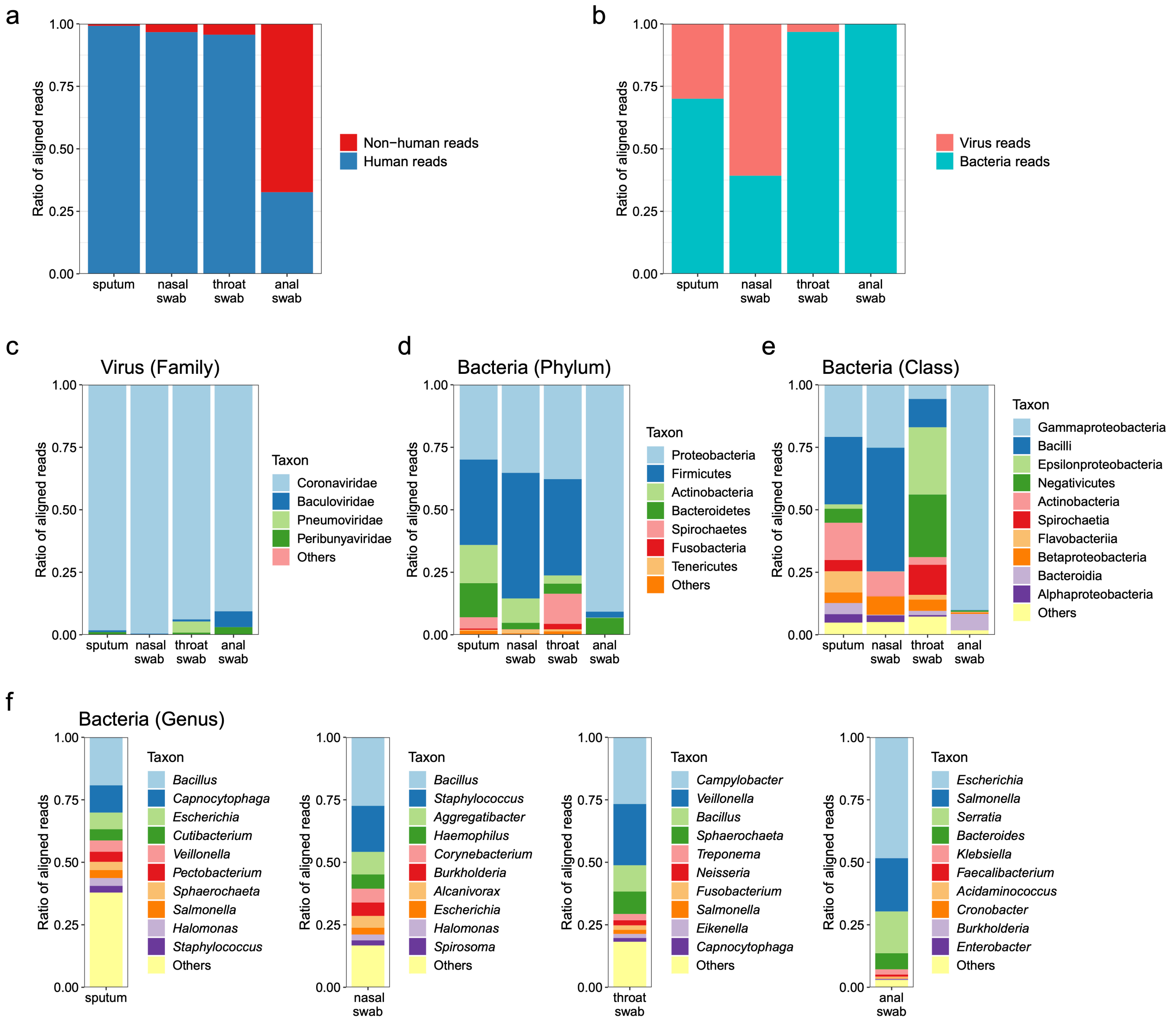


**Supplementary Figure 4. Taxonomy of clinical samples by unbiased metatranscriptomic sequencing.** Distribution of average ratios of human and non-human reads (**a**); virus and bacteria reads (**b**); virus reads at the family level (**c**); and bacteria reads at the phylum (**d**), class (**e**) and genus level (**f**) in metatranscriptomic data of sputum (n=3), nasal swab (n=2), throat swab (n=2) and anal swab (n=1).

**Supplementary table 1.** Capture sequencing data summary of serial dilutions of the cultured isolate and eight HCoV-19 positive clinical samples collected from Guangzhou (PE100).

| Sample ID | Sample type | Ct value | Estimated genome copies per mL | # of sequencing read pairs | # of HCoV-19 read pairs | % of HCoV-19 read pairs | % Coverage | Depth |
|---|---|---|---|---|---|---|---|---|
| **D1** | Cultured isolate | 17.3 | 1.00E+07 | 239,860,837 | 217,024,080 | 90.479 | 100 | 204681.46 |
| **D2** | Cultured isolate | 20.8 | 1.00E+06 | 240,646,932 | 215,939,790 | 89.733 | 100 | 256729.31 |
| **D3** | Cultured isolate | 24.5 | 1.00E+05 | 202,818,485 | 141,387,728 | 69.711 | 100 | 202126.84 |
| **D4** | Cultured isolate | 28.7 | 1.00E+04 | 205,774,008 | 30,282,575 | 14.716 | 99.99 | 42703.91 |
| **D5** | Cultured isolate | 31.8 | 1.00E+03 | 220,321,378 | 1,370,741 | 0.622 | 99.93 | 2117.42 |
| **D6** | Cultured isolate | 35 | 1.00E+02 | 210,213,840 | 67,315 | 0.032 | 99.92 | 99.14 |
| **D7** | Cultured isolate | 39.9 | 1.00E+01 | 202,621,615 | 31,030 | 0.015 | 99.88 | 38.56 |
| **GZMU0047** | nasal swab | 18 | - | 170,047,548 | 155,485,218 | 91.436 | 99.99 | 200186.17 |
| **GZMU0016** | sputum | 21 | - | 156,030,520 | 144,355,833 | 92.518 | 99.99 | 155582.65 |
| **GZMU0048** | throat swab | 24 | - | 237,899,560 | 216,468,337 | 90.991 | 99.91 | 209169.77 |
| **GZMU0044** | nasal swab | 26 | - | 171,428,241 | 158,999,936 | 92.750 | 99.92 | 178822.54 |
| **GZMU0030** | throat swab | 29 | - | 108,293,737 | 44,206,637 | 40.821 | 99.89 | 30352.36 |
| **GZMU0014** | sputum | 30 | - | 106,272,633 | 57,202,971 | 53.827 | 99.9 | 43147.74 |
| **GZMU0042** | sputum | 32 | - | 129,222,354 | 113,772,944 | 88.044 | 99.95 | 107197.11 |
| **GZMU0031** | anal swab | 32 | - | 197,895,055 | 1,162,261 | 0.587 | 99.89 | 747.24 |

**Supplementary table 2.** Amplicon sequencing data summary of serial dilutions of the cultured isolate and eight HCoV-19 positive clinical samples collected from Guangzhou (SE400).

| Sample ID | Sample Type | Ct value | Estimated genome copies per mL | # of sequencing read pairs | # of HCoV-19 read pairs | % of HCoV-19 read pairs | % of HCoV-19 read pairs (-NAB) [a] | % Coverage | Depth |
|---|---|---|---|---|---|---|---|---|---|
| **D2** | Cultured isolate | 20.8 | 1.00E+06 | 3,841,506 | 3,413,916 | 88.87 | 89.04 | 99.79 | 36054.77 |
| **D3** | Cultured isolate | 24.5 | 1.00E+05 | 2,836,864 | 2,454,959 | 86.54 | 88.19 | 99.87 | 25698.05 |
| **D4** | Cultured isolate | 28.7 | 1.00E+04 | 4,640,806 | 3,178,230 | 68.48 | 83.87 | 99.79 | 33132.25 |
| **D5** | Cultured isolate | 31.8 | 1.00E+03 | 10,321,121 | 2,183,454 | 21.16 | 50.38 | 99.79 | 22796.57 |
| **D6** | Cultured isolate | 35 | 1.00E+02 | 10,132,009 | 312,440 | 3.08 | 14.10 | 99.52 | 3242.58 |
| **D7** | Cultured isolate | 39.9 | 1.00E+01 | 8,960,529 | 30,928 | 0.35 | 1.30 | 95.23 | 319.56 |
| **GZMU0047** | nasal swab | 18 | - | 2,327,683 | 2,138,967 | 91.893 | 91.92 | 99.79 | 22215.18 |
| **GZMU0016** | sputum | 21 | - | 4,048,162 | 3,429,988 | 84.730 | 87.85 | 99.79 | 35678.71 |
| **GZMU0048** | throat swab | 24 | - | 2,599,809 | 2,299,379 | 88.444 | 89.77 | 99.79 | 23976.41 |
| **GZMU0044** | nasal swab | 26 | - | 3,104,511 | 2,733,286 | 88.042 | 88.60 | 99.79 | 28560.27 |
| **GZMU0030** | throat swab | 29 | - | 730,663 | 403,509 | 55.225 | 77.81 | 99.79 | 4220.42 |
| **GZMU0014** | sputum | 30 | - | 4,567,919 | 41,233 | 0.903 | 3.46 | 99.79 | 428.82 |
| **GZMU0042** | sputum | 32 | - | 3,877,593 | 160,934 | 4.150 | 17.34 | 99.79 | 1669.91 |
| **GZMU0031** | anal swab | 32 | - | 1,099,212 | 352 | 0.032 | 0.12 | 73.65 | 3.59 |

[a] represents the % of HCoV-19 reads in total reads depleted of the reads from amplified nucleic acids background

**Supplementary Table 3.** Estimated minimum amount of sequencing data required to achieve ≥ 10X sequencing depth and ≥ 95% coverage for different methods.

| Sample ID | Sample Type | Ct value | Estimated genome copies per mL | Meta (bp) | Capture (bp) | Amplicon (bp) |
|---|---|---|---|---|---|---|
| **D0** | Cultured isolate | N.D. | 1.00E+08 | 9,824,200 | N.D. | N.D. |
| **D1** | Cultured isolate | 17.3 | 1.00E+07 | N.D. | 6,384,200 | N.D. |
| **D2** | Cultured isolate | 20.8 | 1.00E+06 | N.D. | 5,313,200 | 33,903,600 |
| **D3** | Cultured isolate | 24.5 | 1.00E+05 | N.D. | 5,655,000 | 34,677,600 |
| **D4** | Cultured isolate | 28.7 | 1.00E+04 | N.D. | 27,228,800 | 43,691,200 |
| **D5** | Cultured isolate | 31.8 | 1.00E+03 | N.D. | 577,615,600 | 141,999,600 |
| **D6** | Cultured isolate | 35 | 1.00E+02 | N.D. | 14,016,023,400 | 973,158,000 |
| **D7** | Cultured isolate | 39.9 | 1.00E+01 | N.D. | 40,524,323,000 (94.19%) [a] | 3,584,211,600 (65.93%) [a] |
| **GZMU0047** | nasal swab | 18 | - | 129,425,800 | 12,562,800 | 32,696,400 |
| **GZMU0016** | sputum | 21 | - | 1,183,209,800 | 15,229,600 | 35,534,800 |
| **GZMU0048** | throat swab | 24 | - | 2,395,109,000 | 16,659,600 | 33,938,400 |
| **GZMU0044** | nasal swab | 26 | - | 1,185,264,800 | 14,565,000 | 34,141,600 |
| **GZMU0030** | throat swab | 29 | - | 243,016,768,200 | 52,231,600 | 54,291,200 |
| **GZMU0014** | sputum | 30 | - | 165,384,913,600 | 37,665,200 | 1,827,167,600 |
| **GZMU0042** | sputum | 32 | - | 11,833,367,600 | 18,421,200 | 723,051,200 |
| **GZMU0031** | anal swab | 32 | - | 334,344,301,400 (56.47%) [a] | 25,844,470,800 (93.99%) [a] | 439,684,800 (8.22%) [a] |

[a] indicates that the total sequencing data of that sample was not sufficient to achieve ≥ 10X sequencing depth and ≥ 95% coverage of HCoV-19 genome. The value within parentheses refers to the percentage of genome covered by ≥ 10X depth using total sequencing data.
N.D. refers to “Not Determined”.

**Supplementary Table 4.** Estimated minimum amount of sequencing data required to achieve high-confidence variants calling analyses for different methods.

| Sample ID | Sample Type | Ct value | Estimated genome copies per mL | Meta | Capture | Amplicon |
|---|---|---|---|---|---|---|
| | | | | (Depth ≥ 10X, CVG ≥ 95%) | (Depth ≥ 20X, CVG ≥ 95%) | (Depth ≥ 100X, CVG ≥ 95%) |
| **D0** | Cultured isolate | N.D. | 1.00E+08 | 9,824,200 | N.D. | N.D. |
| **D1** | Cultured isolate | 17.3 | 1.00E+07 | N.D. | 10,628,200 | N.D. |
| **D2** | Cultured isolate | 20.8 | 1.00E+06 | N.D. | 8,873,800 | 185,765,200 |
| **D3** | Cultured isolate | 24.5 | 1.00E+05 | N.D. | 9,335,000 | 190,712,800 |
| **D4** | Cultured isolate | 28.7 | 1.00E+04 | N.D. | 44,817,400 | 240,931,200 |
| **D5** | Cultured isolate | 31.8 | 1.00E+03 | N.D. | 960,894,400 | 780,488,400 |
| **D6** | Cultured isolate | 35 | 1.00E+02 | N.D. | 24,474,133,200 | 2,757,126,800 |
| **D7** | Cultured isolate | 39.9 | 1.00E+01 | N.D. | 40,524,323,000 (78.8%) [a] | 3,584,211,600 (39.49%) [a] |
| **GZMU0047** | nasal swab | 18 | - | 129,425,800 | 23,701,600 | 261,828,400 |
| **GZMU0016** | sputum | 21 | - | 1,183,209,800 | 28,664,000 | 283,599,200 |
| **GZMU0048** | throat swab | 24 | - | 2,395,109,000 | 31,466,200 | 271,115,200 |
| **GZMU0044** | nasal swab | 26 | - | 1,185,264,800 | 27,208,200 | 271,859,200 |
| **GZMU0030** | throat swab | 29 | - | 243,016,768,200 | 99,263,400 | 292,265,200 (94.15) [a] |
| **GZMU0014** | sputum | 30 | - | 165,384,913,600 | 70,173,000 | 1,827,167,600 (71.66%) [a] |
| **GZMU0042** | sputum | 32 | - | 11,833,367,600 | 34,466,600 | 1,551,037,200 (88.17%) [a] |
| **GZMU0031** | anal swab | 32 | - | 334,344,301,400 (56.47%) [a] | 25,844,470,800 (67.06%) [a] | 439,684,800 (0.00%) [a] |

[a] indicates that the total sequencing data of a given sample was not sufficient to achieve method-specific requirement. The value within parentheses refers to the percentage of genome covered by ≥ method-specific depth using total sequencing data.
N.D. refers to “Not Determined

**Supplementary Table 5.** Genome positions of amplicons. (see SupplementaryTable5.xlsx)

**Supplementary Table 6.** Bioinformatic tools and parameters used in this study.

**1. Identification of HCoV-19-like reads from Massively Parallel Sequencing data**

| Software | Version | Parameters |
|---|---|---|
| Kraken | v0.10.5 | dafault |
| fastp | v0.19.5 | -q 20 -u 20 -n 1 -l 50 |
| SOAPnuke | v1.5.6 | -l 20 -q 0.2 -E 50 -n 0.02 -5 0 -Q 2 -G -d |
| PRINSEQ | v0.20.4 | -lc_method dust -lc_threshold 7 |
| fastp | v0.19.5 | -q 20 -u 20 -n 1 -l 50 |
| BWA aln | v0.7.16 | default |
| BAMClipper | v1.1.1 | -n 4 -u 21 -d 21 |

**2. Assembling viral genome**

| Software | Version | Parameters |
|---|---|---|
| SPAdes | v3.14.0 | default |
| Pilon | v1.23 | --changes –vcf --changes --vcf --mindepth 1 --fix all, amb |

**3.Assessment the coverage depth across the viral genome**

| Software | Version | Parameters |
|---|---|---|
| Picard | v2.10.10 | Markduplicates |
| Samtools | v1.9 | depth |
| R | v3.6.1 | ggplot2 |
| R | v3.6.1 | pheatmap |

**4.Relationships between genome copies and method-dependent minimum amount of sequencing data**

| Software | Version | Parameters |
|---|---|---|
| Samtools | v1.9 | view -s |
| R | v3.6.1 | ggplot2 |
| R | v3.6.1 | ggscatter |

**5.Consistency in variants calling performance among methods**

| Software | Version | Parameters |
|---|---|---|
| freebayes | v1.3.1 | -p 1 -q 20 -m 60 --min-coverage 10 -V |
| snippy-vcf_filter | v3.2 | --minqual 100 --mincov 10 --minfrac 0.8 |
| SNPeff | v4.3 | default |
| pysamstats | v1.1.2 | -type variation_strand --min-baseq 20 -D 1000000 |

**6.Taxonomy of clinical samples by unbiased metatranscriptomic sequencing**

| Software | Version | Parameters |
|---|---|---|
| fastp | v0.19.5 | -5 -3 -q 20 -c -l 30 |
| SOAP2 | v2.21 | -m 0 -x 1000 -s 28 -l 32 -v 5 -r 1 |
| HISAT2 | v2.1.0 | default |
| Kraken 2 | v2.0.8-beta | --threads 24 --confidence 0 |
| Bracken | v2.5.0 | default |

**Supplementary Table 7.** Genome sequences acknowledged. (see SupplementaryTable7.xlsx)
